# The complete mitochondrial genome of Spurilla braziliana MacFarland 1909 (Nudibranchia, Aeolidiidae)

**DOI:** 10.1101/2023.04.04.535430

**Authors:** Hideaki Mizobata, Kentaro Hayashi, Ryo Yonezawa, Andre Lanza, Shigeharu Kinoshita, Kazutoshi Yoshitake, Shuichi Asakawa

## Abstract

*Spurilla braziliana* MacFarland 1909 is a morphologically diverse nudibranch found in the Pacific and Western Atlantic. The complete mitochondrial genome of *S. braziliana* has been constructed using next-generation sequencing technology. The mitochondrial genome is 14,291 bp and contains 13 protein-coding genes, 2 rRNA genes, and 23 tRNA genes. Molecular phylogenetic analysis using the maximum likelihood method revealed that *S. braziliana* is included in the superfamily Aeolidioidea and forms a monophyletic group with *Berghia stephanieae*, a nudibranch of the family Aeolidiidae. This study reinforces existing taxonomic insights and provides a basis for further molecular phylogenetic analysis.

## Introduction

Members of the genus *Spurilla* are unique nudibranchs capable of storing and utilizing cnidarian nematocysts in their bodies (Greenwood and Mariscal 1984). These sea slugs are widely distributed in the world’s oceans and attract attention from a taxonomic perspective due to their diverse coloration and morphological characteristics (Carmona et al. 2014). Previous research by Carmona et al. (Carmona et al. 2013) has shown that only one species of *Spurilla, S. braziliana*, is found in the Pacific Ocean. However, there is significant morphological diversity within *S. braziliana* (Carmona et al. 2014) suggesting that further molecular phylogenetic analysis may be required to distinguish the different morphotypes. This study reports the complete mitochondrial genome sequence of one morphotype of *S. braziliana* from Japan.

## Materials

A sample of *Spurilla braziliana* was collected on June 10, 2021 from the intertidal area of Miura City (35°16’18.8”N 139°34’04.7”E), Kanagawa Prefecture, Japan (Figure 1) and preserved in NucleoProtect RNA (MACHEREY-NAGEL) for 5.5 months at -80 °C before DNA extraction. The sample was then transferred to 99% ethanol for deposition at the University Museum of the University of Tokyo (http://www.um.u-tokyo.ac.jp/, Assoc. Prof. Takenori Sasaki) under voucher number RM34044.

**Figure 1.**
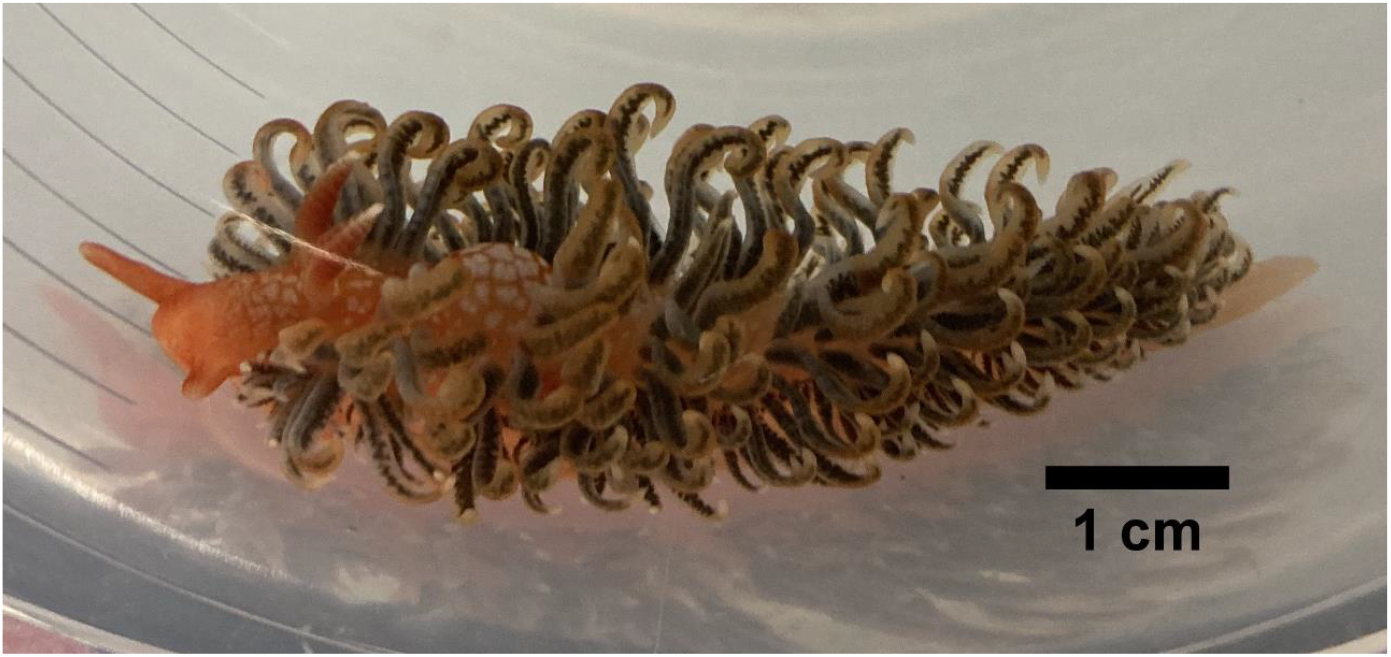
The individual of *S. braziliana* used in this study. Its morphotype is similar to the one shown in Fig. 4E in the previous taxonomic study (Carmona et al. 2014). The picture was taken by Hideaki Mizobata.

## Methods

Prior to the sequencing process, genomic DNA was extracted from 25 mg of the body wall of the aforementioned sample using the DNeasy Blood & Tissue Kit (QIAGEN) according to the manufacturer’s instructions. A DNAlibrary for PE150 sequencing was prepared with MGIEasy FS DNA Library Prep Set (MGI) and sequenced with DNBSEQ-T7 (MGI) at Genome-Lead Corporation, Japan.A total of 63.6 Gbp of sequence data was obtained from the sequencing, with a Q30 score of 94.08%, indicating sufficient quality for assembly. The reads were assembled with CLC Genomics Workbench version 8.5 (QIAGEN Aarhus A/S) to construct a primary assembly. The primary assembly was polished using bwa mem version 0.7.15-r1140 (Li 2013), samtools version 1.4 (Danecek et al. 2021), and pilon version 1.22 (Walker et al. 2014) with the following commands.

$ bwa index assembly.fasta

$ bwa mem assembly.fasta read1.fastq read2.fastq|samtools view -bS|samtools sort > assembly.fasta.bam

$ samtools index assembly.fasta.bam

$ pilon –genome assembly.fasta –frags assembly.fasta.bam –outdir.

To further improve the polished primary assembly, the sequence was circularized by the ends, cut at a site differing from the circularization point, and polished once more. The raw reads were mapped to the final mitochondrial genome using bwa mem and samtools to ensure proper assembly (Supplementary Figure 1). The average coverage was confirmed to be 7866, further validating the robustness of the assembly and polishing. The polished mitochondrial genome was annotated with MITOS (Donath et al. 2019) and manually corrected using Geneious Prime Java version 11.0.14.1+1 (Biomatters Ltd.). MitoZ (Meng et al. 2019) was used for the accurate annotation of tRNA. The identified tRNA genes were initially folded using the RNA fold web server (Gruber et al. 2008) to generate a rough secondary structure, which was then manually adjusted using VARNA (Darty et al. 2009). Annotations and other mitochondrial genome structures were visualized using OGDRAW (Greiner et al. 2019).

To confirm the species identity of our sample as *S. braziliana*, we performed nucleotide BLAST (blastn) analyses using the web interface of the National Center for Biotechnology Information (NCBI). The *COX1* and *16S rRNA* gene sequences from our specimen were used as queries, with the non-redundant nucleotide collection (nt) database chosen for the search. Default parameters were used for all other settings.

The 13 protein-coding genes within the mitochondrial genome were each individually aligned with their corresponding genes from 22 known Nudibranchia species’ mitochondrial genomes (Table 1) using MAFFT v7.508 (Katoh and Standley 2013). This was done based on their amino acid sequences, as illustrated in Supplementary Data 1. For each alignment, 7 species belonging to the suborder Cladobranchia were selected. The selected 13 alignments were then concatenated. Based on the concatenated alignment, the molecular phylogenetic tree was generated with the maximum likelihood method with 1000 bootstrap replicates in MEGA version 10.1.8 (Kumar et al. 2018).

**Table 1.** The list of species, GenBank accession ID, and references used for alignments (Supplementary Data 1). Species used to create the molecular phylogenetic tree (Fig. 3) are indicated by an asterisk.

| Scientific name | GenBank accession ID | Reference |
| --- | --- | --- |
| * *Spurilla braziliana* | LC759638 | Present study |
| *Goniobranchus leopardus* | MZ747465 | - |
| *Chromodoris orientalis* | MH550543 | Yu et al. 2018 |
| *Glossodoris acosti* | MZ823409 | - |
| *Hypselodoris festiva* | KU365323 | Karagozlu et al. 2016 |
| *Verconia nivalis* | OL800586 | Do et al. 2022 |
| *Polycera hedgpethi* | MZ713367 | - |
| *Triopha modesta* | MW387958 | Do et al. 2022 |
| *Doris odhneri* | OL800585 | Do et al. 2022 |
| *Asteronotus hepaticus* | MW559976 | - |
| *Carminodoris armata* | OL800584 | Do et al. 2022 |
| *Notodoris gardineri* | DQ991934 | Medina et al. 2011 |
| *Phyllidiopsis krempfi* | MT726194 | Kim et al. 2021 |
| *Phyllidiella pustulosa* | MK279705 | Do et al. 2019 |
| *Roboastra europaea* | AY083457 | Grande et al. 2004 |
| *Phestilla sp.* | PN553001 | - |
| * *Dermatobranchus otome* | MT527185 | Do et al. 2020 |
| * *Tritonia diomedea* | KP764765 | Sevigny et al. 2015 |
| * *Melibe leonina* | KP764764 | Sevigny et al. 2015 |
| * *Sakuraeolis japonica* | KX610997 | Karagozlu et al. 2016 |
| * *Hermisenda emurai* | MK279704 | Do et al. 2019 |
| * *Berghia stephanieae* | MW027646 | Clavijo et al. 2021 |

## Results and Discussion

The mitochondrial genome of *S. braziliana* (GenBank Accession ID: LC759638) is 14,291 bp and has a GC content of 33.94%. It consists of 13 protein-coding genes, 2 rRNA genes, and 23 tRNA genes (Figure 2). All 13 protein-coding genes exhibit high amino acid sequence similarity with those of other nudibranch species (Supplementary Data 1), suggesting the accuracy of the genome annotation. The gene order is identical to that of *Sakuraeolis japonica* (Karagozlu et al. 2016) and *Hermissenda emurai* (Do et al. 2019), both of which belong to the superfamily Aeolidioidea, the same as *S. braziliana*. The start codons for *S. braziliana* were compared with those of 22 reference species (with 21 species for *ND5* gene). This comparison revealed six different start codons in 23 nudibranchs: ATA, ATT, ATG, ATC, TTG, GTG. The start codon usage in *S. braziliana* was found to be distinct, particularly the use of GTG in *ATP6, ATP8, COX1*, and *COX2* genes, a pattern rarely observed in the comparison species (*ATP6*: 1/23 species, *ATP8*: 1/23, *COX1*: 6/23, *COX2*: 1/23). Within the NADH dehydrogenase gene series, TTG serves as the start codon in 5 genes, excluding *ND4* and *ND5*. For the remaining 4 genes, *COX3, CYTB*, and *ND4* adopt ATG as their start codon, while *ND5* employs ATA. *Berghia stephanieae*, which belongs to the same family, Aeolidiidae, as *S. braziliana*, has been reported to exhibit *tRNA-Ser 1* gene duplication (Clavijo et al. 2021). In our *S. braziliana* sample as well, gene duplications of *tRNA-Ser* were observed, and these tRNAs exhibited a unique secondary structure with a missing D-arm (Supplementary Data 2).

**Figure 2.**
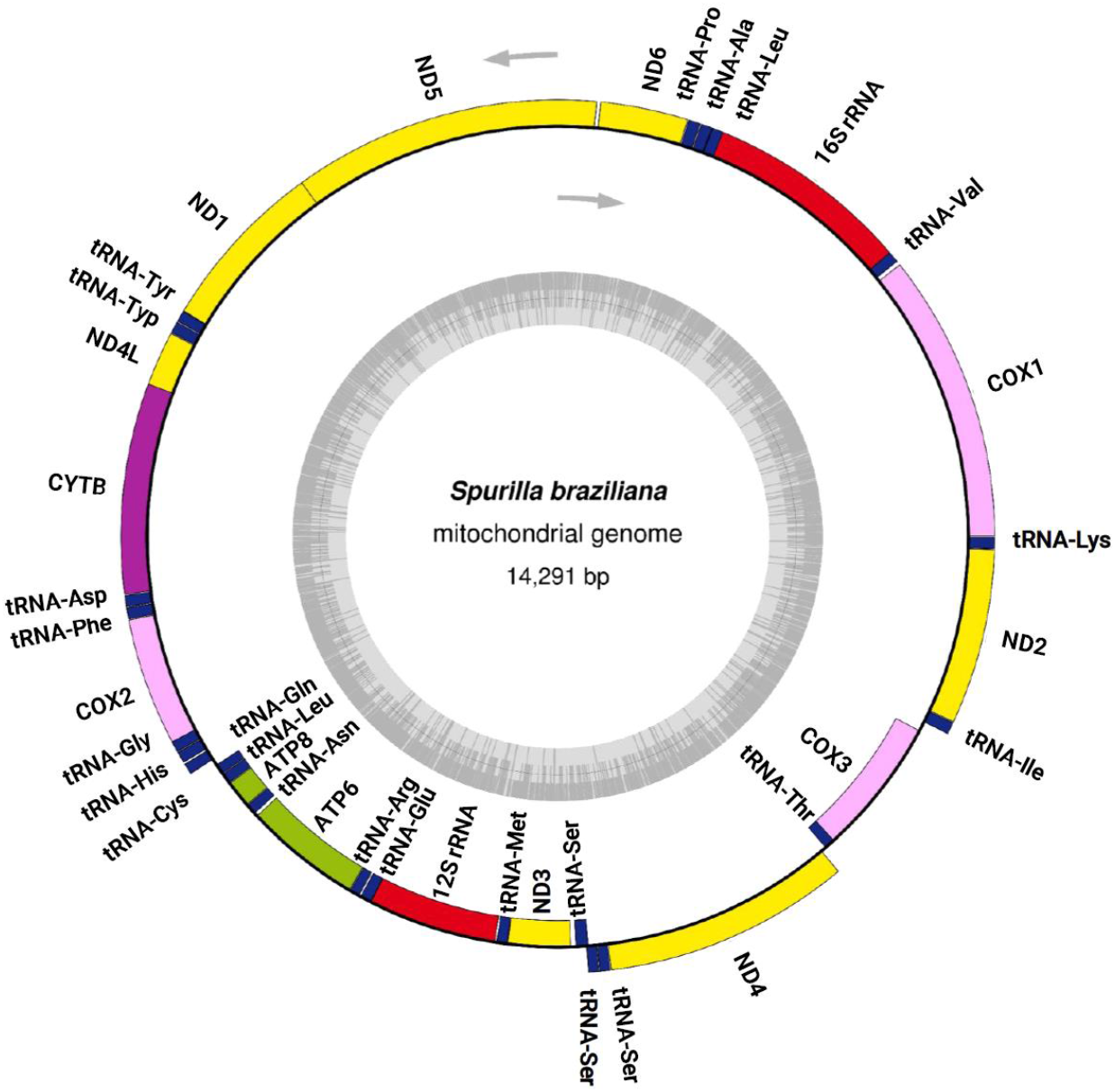
Circular map of the complete mitochondrial genome annotated to show the locations and orientation of its 13 protein-coding genes, 2 rRNA genes, and 23 tRNA genes. The upper arrows indicate the direction of transcription, while the inner gray circle represents the GC content.

Through our blastn analyses of the annotated *COX1* and *16S rRNA* genes, we found that they only exhibited over 90% similarity with two species, *S. braziliana* and *S. neapolitana*, thus confirming that our sample belongs to the genus *Spurilla*. The *COX1* gene showed 100% similarity with one of the *S. braziliana* references, but did not demonstrate complete similarity with the *S. neapolitana* reference. Given that all *Spurilla* nudibranchs found in the Pacific are *S. braziliana* (Carmona et al., 2013), we can confidently assert that our specimen is *S. braziliana*.

As shown in the molecular phylogenetic tree (Figure 3), *S. braziliana* forms a monophyletic group with *B. stephanieae* and members of the superfamily Aeolidioidea..

**Figure 3.**
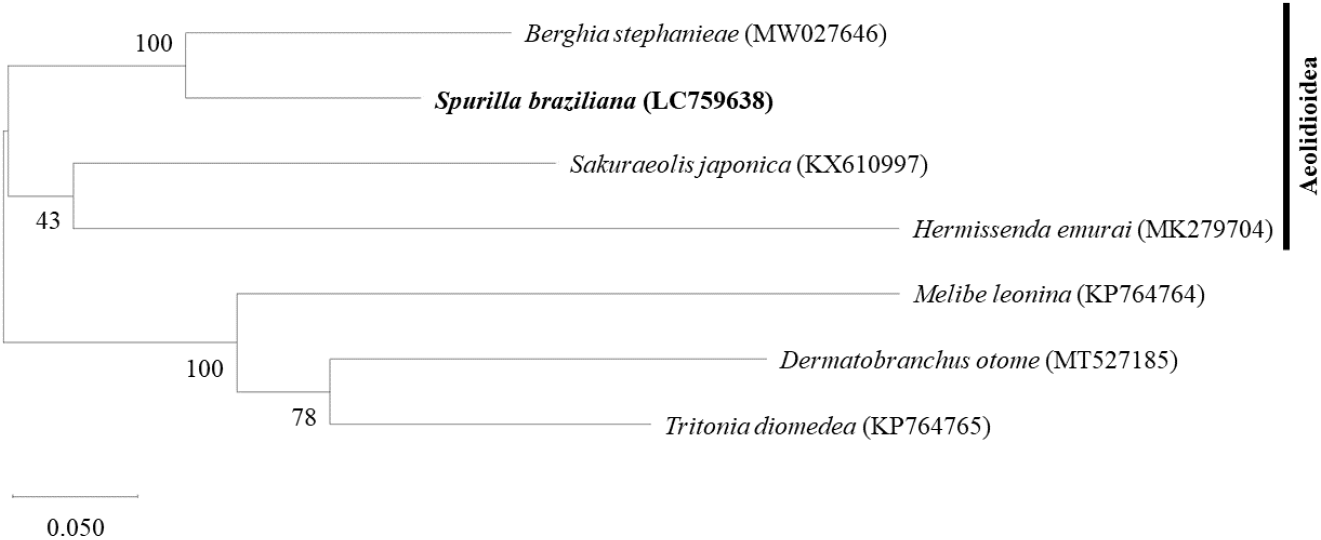
Maximum likelihood phylogenetic tree of *S. braziliana* and 6 other Cladobranchia species based on the concatenated amino acid alignments of the 13 protein-coding genes in the mitochondrial genome. Bootstrap support values are shown for each node, based on 1000 replicates. The list of species and their GenBank accession IDs are shown in Table 1.

The previous molecular phylogenetic study of the superfamily Aeolidioidea using *COI, 16S rRNA*, and *H3* genes showed that the monophyletic group containing the genera *Spurilla* and *Berghia* was a sister group to the group containing *Sakuraeolis* (Carmona et al. 2011). The phylogenetic tree generated using all the protein-coding genes of the mitochondrial genome matched these findings. The mitochondrial genome of *S. braziliana*, the first mitochondrial genome of its genus, will provide important data for future taxonomic studies on *Spurilla* nudibranchs, which are known for their highly variable morphology.

## Ethical approval

The “Manual for Animal Experiment of the University of Tokyo” provides ethical guidelines only for mammals, birds, and reptiles, and does not require ethical review for the invertebrates used in this study. Therefore, any specific permission from the “Office for Life Science Research Ethics and Safety” of the University of Tokyo is not necessary for this research.

## Author contributions

H.M., R.Y., and K.Y. formulated the research concept. H.M., K.H., and R.Y. collected the *S. braziliana* samples. H.M. conducted the experimental manipulation and analysis, excluding library preparation and next-generation sequencing tasks. Supervision was provided by S.K., K.Y., and S.A. All authors contributed to data interpretation and discussion of the results. The manuscript was written by H.M. and A.L. All authors participated in the revision of the manuscript draft and approved the final version. All authors agree to be accountable for all aspects of the work and ensure that inquiries concerning the accuracy or integrity of any part of the work are thoroughly investigated and resolved.

## Data availability

The genome sequence data that support the findings of this study are openly available in GenBank of NCBI at https://www.ncbi.nlm.nih.gov under the accession ID: LC759638. The associated BioProject, SRA, and Bio-sample numbers are PRJDB15401, DRR450785, and SAMD00585630, respectively.

---

**Supplementary Figure 1.**
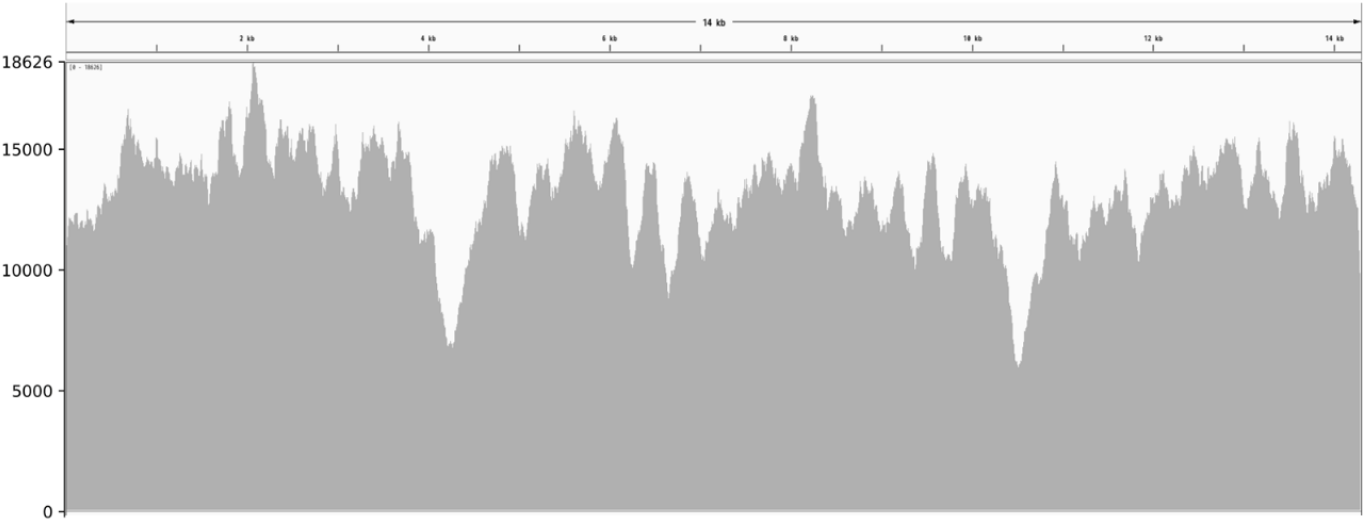
Read-depth plot for the mitochondrial genome.

**Supplementary Data 1.**
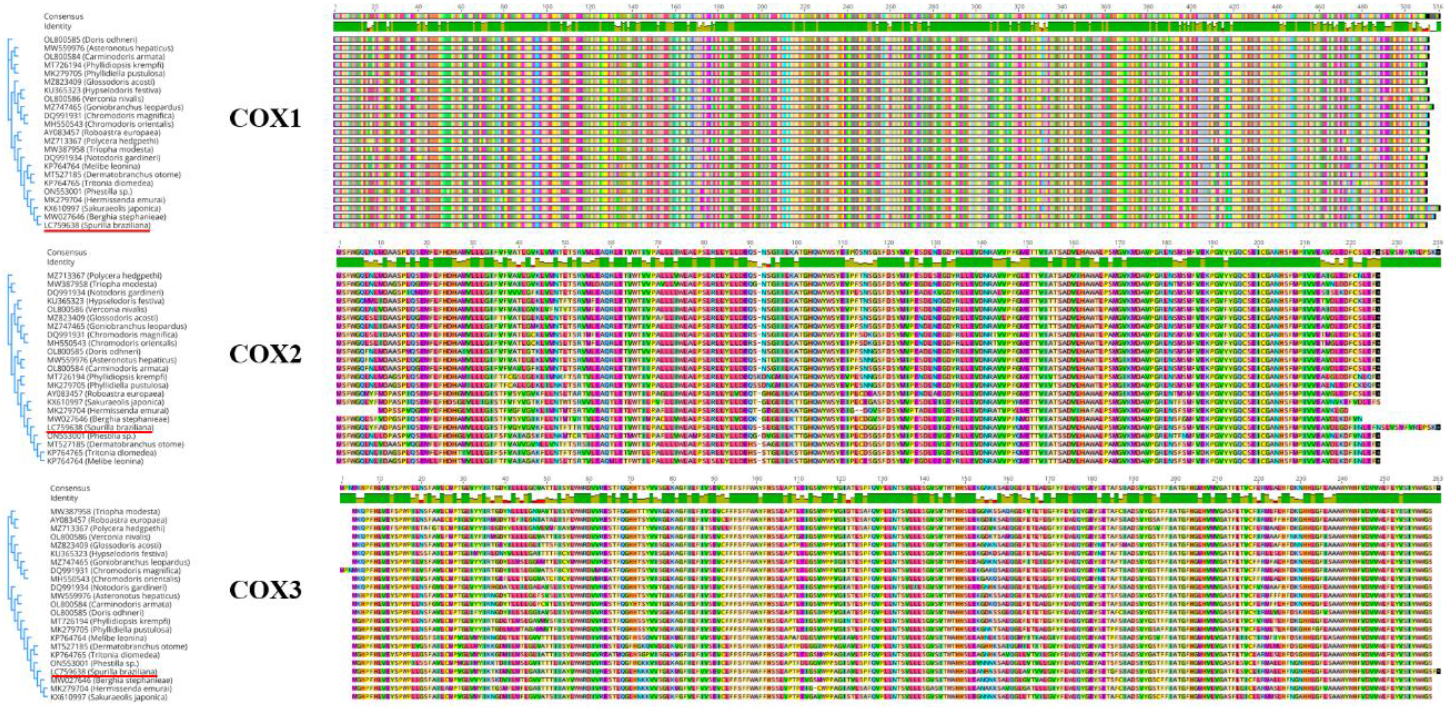
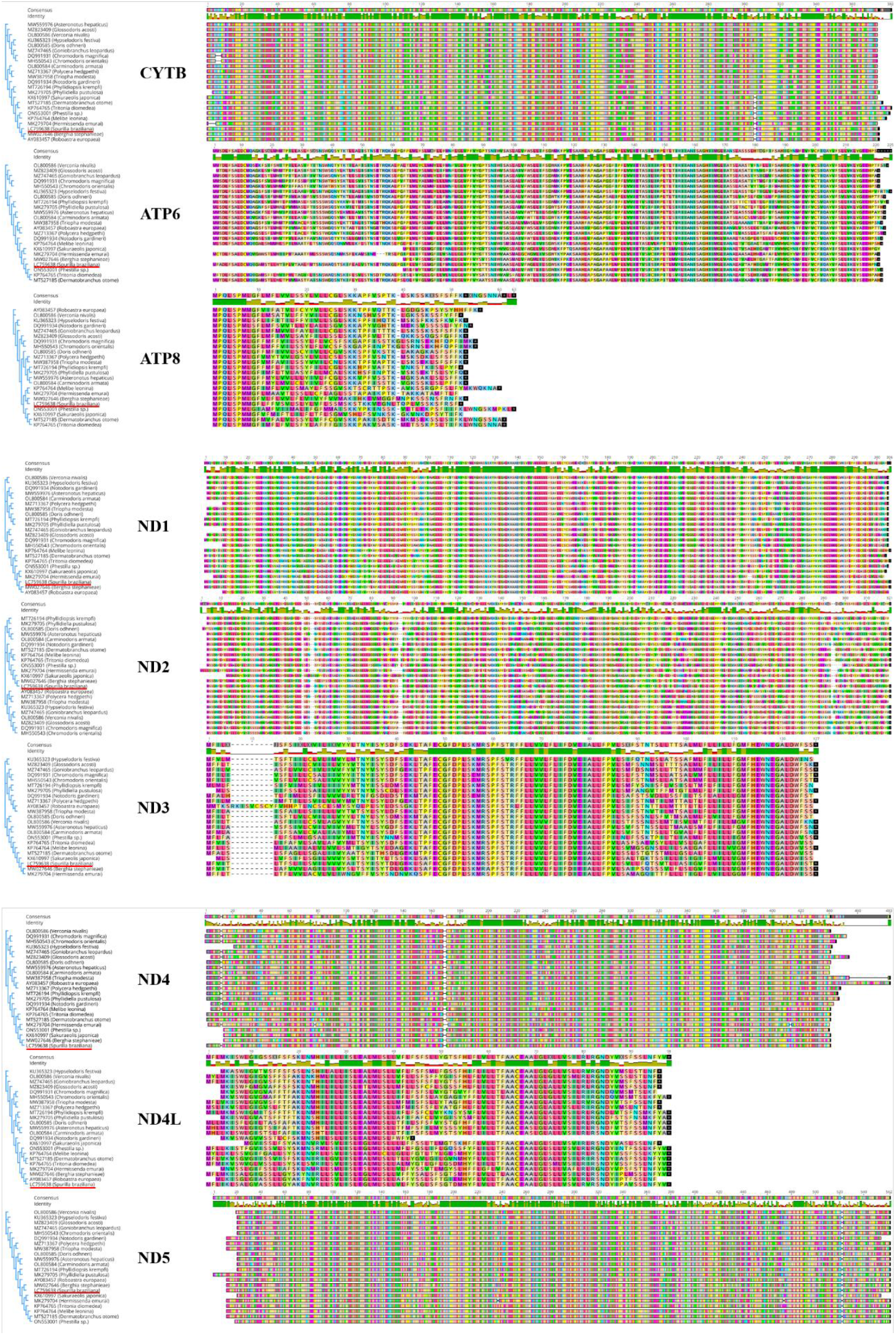
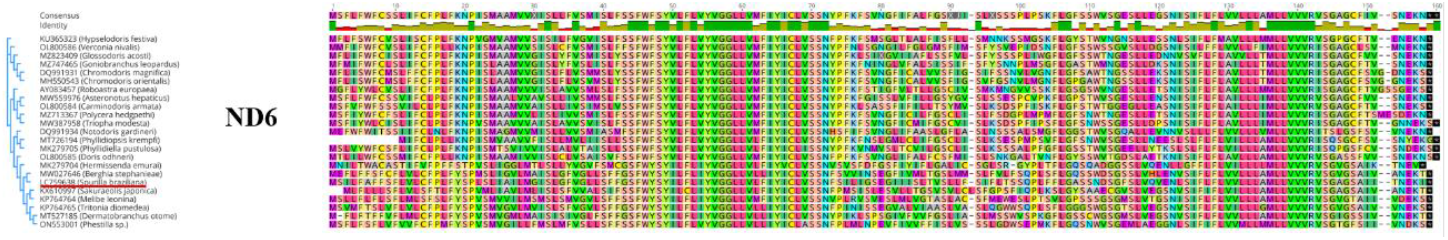
Alignments for 13 protein-coding genes across the mitochondrial genome of *Spurilla braziliana* and those of 22 previously characterized nudibranch species.

**Supplementary Data 2.**
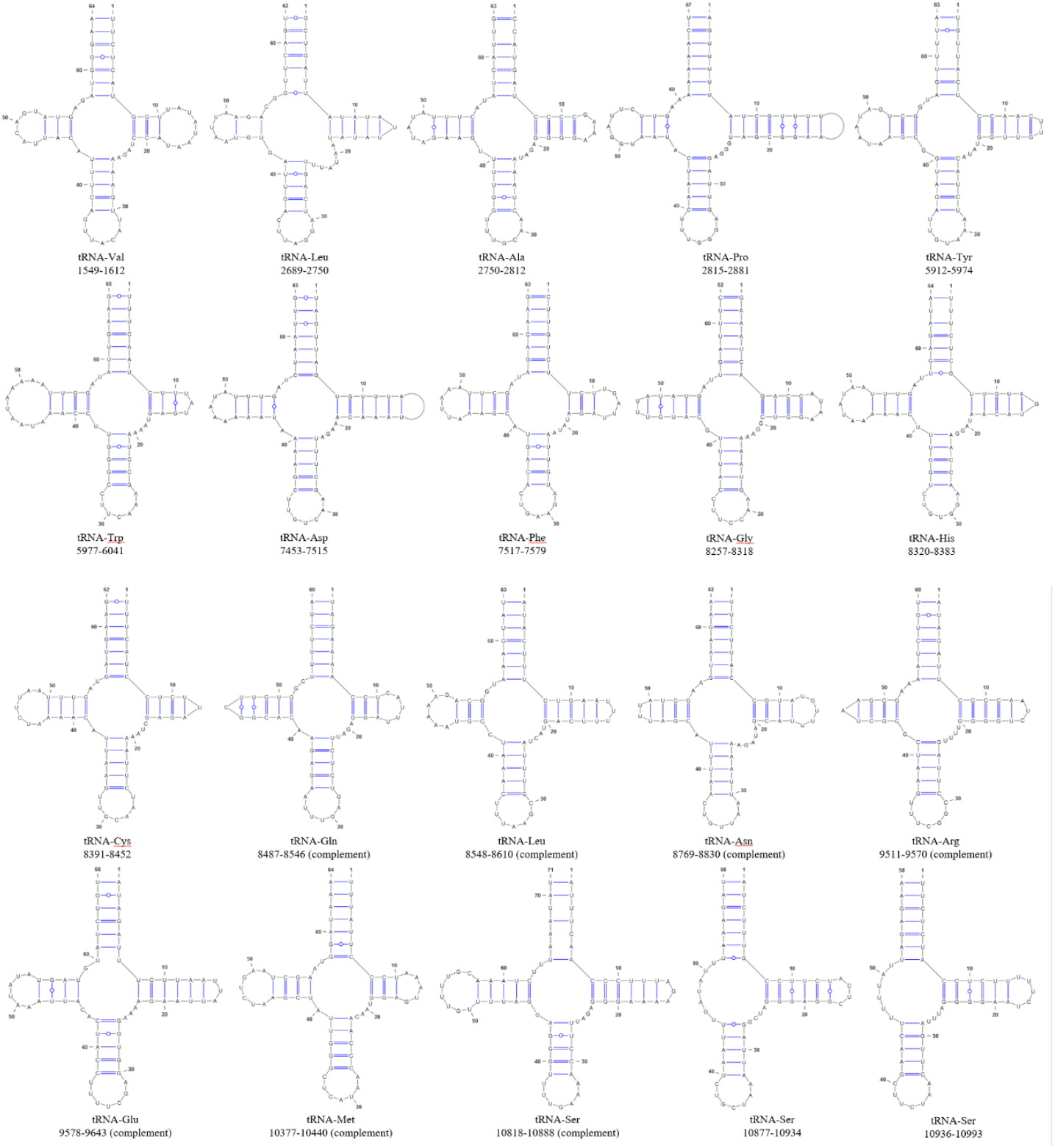
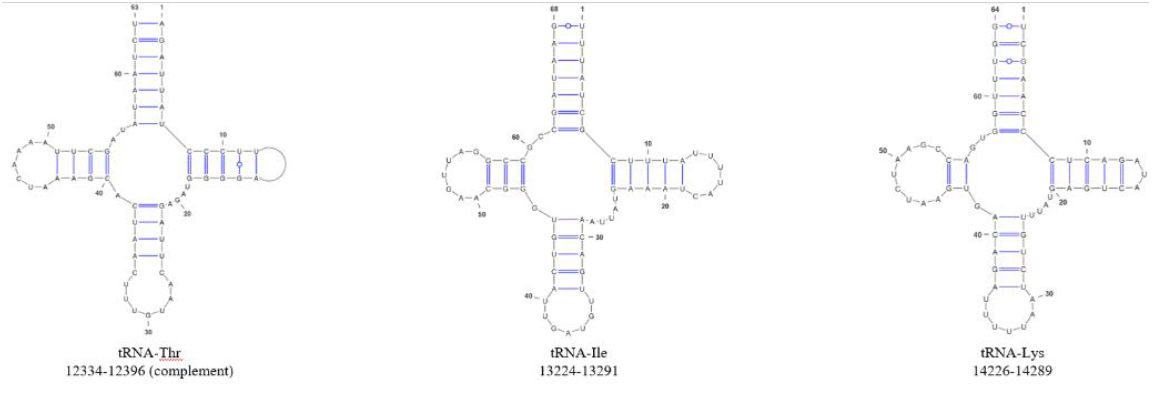
The secondary structures of the 23 tRNAs encoded in the mitochondrial genome of *Spurilla braziliana*. The types of base pairing are indicated by the varying styles of bonds between bases. Each structure is accompanied by the corresponding tRNA type and its position within the mtDNA.

## References

Carmona, Leila, Terrence M. Gosliner, Marta Pola, and Juan Lucas Cervera. 2011. ‘A Molecular Approach to the Phylogenetic Status of the Aeolid Genus Babakina Roller, 1973 (Nudibranchia)’. Journal of Molluscan Studies 77 (4): 417–422. doi:10.1093/mollus/eyr029.

Carmona, Leila, Bonnie R. Lei, Marta Pola, Terrence M. Gosliner, Angel Valdes, and Cervera. 2014. ‘Untangling the Spurilla Neapolitana (Delle Chiaje, 1841) Species Complex: A Review of the Genus Spurilla Bergh, 1864 (Mollusca: Nudibranchia: Aeolidiidae)’. Zoological Journal of the Linnean Society, January. doi:10.1111/zoj12098.

Carmona, Leila, Marta Pola, Terrence M. Gosliner, and Juan Lucas Cervera. 2013. ‘A Tale That Morphology Fails to Tell: A Molecular Phylogeny of Aeolidiidae (Aeolidida, Nudibranchia, Gastropoda)’. Edited by Jonathan H. Badger. PLoS ONE 8 (5): e63000. doi:10.1371/journal.pone.0063000.

Danecek, Petr, James K Bonfield, Jennifer Liddle, John Marshall, Valeriu Ohan, Martin O Pollard, Andrew Whitwham, et al. 2021. ‘Twelve Years of SAMtools and BCFtools’. GigaScience 10 (2): giab008. doi:10.1093/gigascience/giab008.

Darty, Kévin, Alain Denise, and Yann Ponty. 2009. ‘VARNA: Interactive Drawing and Editing of the RNA Secondary Structure’. Bioinformatics 25 (15): 1974–1975. doi:10.1093/bioinformatics/btp250.

Dinh Do, Thinh Tae-June Choi, Dae-Wui Jung, Jung-Il Kim, Mustafa Zafer Karagozlu, and Chang-Bae Kim. 2019. ‘The Complete Mitochondrial Genome of Phyllidiella Pustulosa (Cuvier, 1804) (Nudibranchia, Phyllidiidae)’. Mitochondrial DNA Part B 4 (1): 771–772. doi:10.1080/23802359.2019.1565976.

Dinh Do, Thinh, Jung-Il Kim, Dae-Wui Jung, Tae-June Choi, Mustafa Zafer Karagozlu, and Chang-Bae Kim. 2019. ‘Characterization of the Complete Mitochondrial Genome of Hermissenda Emurai (Baba, 1937) (Nudibranchia, Facelinidae)’. Mitochondrial DNA Part B 4 (1): 860–861. doi:10.1080/23802359.2019.1572477.

Do, Thinh Dinh, Yisoo Choi, Dae-Wui Jung, and Chang-Bae Kim. 2020. ‘Caution and Curation for Complete Mitochondrial Genome from Next-Generation Sequencing: A Case Study from Dermatobranchus Otome (Gastropoda, Nudibranchia)’. Animal Systematics, Evolution and Diversity 36 (4): 336–346. doi:10.5635/ASED.2020.36.4.039.

Do, Thinh Dinh, Dae-Wui Jung, and Chang-Bae Kim. 2022. ‘Molecular Phylogeny of Selected Dorid Nudibranchs Based on Complete Mitochondrial Genome’. Scientific Reports 12 (1): 18797. doi:10.1038/s41598-022-23400-9.

Donath, Alexander, Frank Jühling, Marwa Al-Arab, Stephan H Bernhart, Franziska Reinhardt, Peter F Stadler, Martin Middendorf, and Matthias Bernt. 2019. ‘Improved Annotation of Protein-Coding Genes Boundaries in Metazoan Mitochondrial Genomes’. Nucleic Acids Research 47 (20): 10543–10552. doi:10.1093/nar/gkz833.

Grande, Cristina, Josè Templado, J. Lucas Cervera, and Rafael Zardoya. 2004. ‘Phylogenetic Relationships among Opisthobranchia (Mollusca: Gastropoda) Based on Mitochondrial Cox 1, TrnV, and RrnL Genes’. Molecular Phylogenetics and Evolution 33 (2): 378–388. doi:10.1016/j.ympev.2004.06.008.

Greenwood, Paul G., and Richard N. Mariscal. 1984. ‘The Utilization of Cnidarian Nematocysts by Aeolid Nudibranchs: Nematocyst Maintenance and Release in Spurilla’. Tissue and Cell 16 (5): 719–730. doi:10.1016/0040-8166(84)90005-3.

Greiner, Stephan, Pascal Lehwark, and Ralph Bock. 2019. ‘OrganellarGenomeDRAW (OGDRAW) version 1.3.1: Expanded Toolkit for the Graphical Visualization of Organellar Genomes’. Nucleic Acids Research 47 (W1): W59–W64. doi:10.1093/nar/gkz238.

Gruber, A. R., R. Lorenz, S. H. Bernhart, R. Neubock, and I. L. Hofacker. 2008. ‘The Vienna RNA Websuite’. Nucleic Acids Research 36 (Web Server): W70–W74. doi:10.1093/nar/gkn188.

Karagozlu, Mustafa Zafer, Jin-Mo Sung, JeaHyun Lee, Seong-Geun Kim, and Chang-Bae Kim. 2016. ‘Complete Mitochondrial Genome Analysis of Sakuraeolis Japonica (Baba, 1937) (Mollusca, Gastropoda, Nudibranchia)’. Mitochondrial DNA Part B 1 (1): 720–721. doi:10.1080/23802359.2016.1229587.

Karagozlu, Mustafa Zafer, JinMo Sung, JeaHyun Lee, Woori Kwak, and Chang-Bae Kim. 2016. ‘Complete Sequences of Mitochondrial Genome of Hypselodoris Festiva (A. Adams, 1861) (Mollusca, Gastropoda, Nudibranchia)’. Mitochondrial DNA Part B 1 (1): 266–267. doi:10.1080/23802359.2016.1159933.

Katoh, K., and D. M. Standley. 2013. ‘MAFFT Multiple Sequence Alignment Software Version 7: Improvements in Performance and Usability’. Molecular Biology and Evolution 30 (4): 772–780. doi:10.1093/molbev/mst010.

Kim, Hana, Moonguen Yoon, Keun-Yong Kim, and Yun-Hwan Jung. 2021. ‘The Complete Mitochondrial Genome of Sea Slug Phyllidiopsis Krempfi Pruvot-Fol, 1957 (Nudibranchia, Phyllidiidae) from Pacific Ocean’. Mitochondrial DNA Part B 6 (4): 1523–1524. doi:10.1080/23802359.2020.1823898.

Kumar, Sudhir, Glen Stecher, Michael Li, Christina Knyaz, and Koichiro Tamura. 2018. ‘MEGA X: Molecular Evolutionary Genetics Analysis across Computing Platforms’. Edited by Fabia Ursula Battistuzzi. Molecular Biology and Evolution 35 (6): 1547–1549. doi:10.1093/molbev/msy096.

Li, Heng. 2013. ‘Aligning Sequence Reads, Clone Sequences and Assembly Contigs with BWA-MEM’. arXiv. doi:10.48550/ARXIV.1303.3997.

Medina, Mónica, Shruti Lal, Yvonne Vallès, Tori L. Takaoka, Benoît A. Dayrat, Jeffrey L. Boore, and Terrence Gosliner. 2011. ‘Crawling through Time: Transition of Snails to Slugs Dating Back to the Paleozoic, Based on Mitochondrial Phylogenomics’. Marine Genomics 4 (1): 51–59. doi:10.1016/j.margen.2010.12.006.

Melo Clavijo, Jenny Franziska Drews, Marcello Pirritano, Martin Simon, Abdulrahman Salhab, Alexander Donath, Silja Frankenbach, et al. 2021. ‘The Complete Mitochondrial Genome of the Photosymbiotic Sea Slug Berghia Stephanieae (Valdés, 2005) (Gastropoda, Nudibranchia)’. Mitochondrial DNA Part B 6 (8): 2281–2284. doi:10.1080/23802359.2021.1914211.

Meng, Guanliang, Yiyuan Li, Chentao Yang, and Shanlin Liu. 2019. ‘MitoZ: A Toolkit for Animal Mitochondrial Genome Assembly, Annotation and Visualization’. Nucleic Acids Research 47 (11): e63–e63. doi:10.1093/nar/gkz173.

Sevigny, Joseph L., Lauren E. Kirouac, William Kelley Thomas, Jordan S. Ramsdell, Kayla E. Lawlor, Osman Sharifi, Simarvir Grewal, et al. 2015. ‘The Mitochondrial Genomes of the Nudibranch Mollusks, Melibe Leonina and Tritonia Diomedea, and Their Impact on Gastropod Phylogeny’. Edited by Bi-Song Yue. PLOS ONE 10 (5): e0127519. doi:10.1371/journal.pone.0127519.

Walker, Bruce J., Thomas Abeel, Terrance Shea, Margaret Priest, Amr Abouelliel, Sharadha Sakthikumar, Christina A. Cuomo, et al. 2014. ‘Pilon: An Integrated Tool for Comprehensive Microbial Variant Detection and Genome Assembly Improvement’. Edited by Junwen Wang. PLoS ONE 9 (11): e112963. doi:10.1371/journal.pone.0112963.

Yu, Cheol, Hana Kim, Hyung June Kim, and Yun-Hwan Jung. 2018. ‘The Complete Mitochondrial Genome of the Oriental Sea Slug: Chromodoris Orientalis (Nudibranchia, Chromodorididae)’. Mitochondrial DNA Part B 3 (2): 1017–1018. doi:10.1080/23802359.2018.1508381.

